# How predictability affects habituation to novelty?

**DOI:** 10.1101/2020.07.24.219253

**Authors:** Kazutaka Ueda, Takahiro Sekoguchi, Hideyoshi Yanagisawa

## Abstract

One becomes accustomed to repeated exposures, even for a novel event. In the present study, we investigated how predictability affects habituation to novelty by applying a mathematical model of arousal that we previously developed, and conducted a psychophysiological experiment to test the model prediction. We formalized habituation to novelty as a decrement in Kullback-Leibler divergence from Bayesian prior to posterior (i.e., information gain) representing arousal evoked from a novel event through Bayesian update. The model predicted an interaction effect between initial uncertainty and initial prediction error (i.e., predictability) on habituation to novelty: The greater the initial uncertainty, the faster the information gain decreases (i.e., the sooner one is habituated). Experimental results using subjective reports of surprise and event-related potential (P300) evoked by visual-auditory incongruity supported the model prediction. Our findings suggest that in highly uncertain situations, repeated exposure to stimuli may enhance habituation to novel stimuli.

## 1 Introduction

Novelty is an essential attribute of creativity. Berlyne examined the relationship between novelty and emotion and stated that if familiarity is too high, people will not feel comfortable; however, if novelty is too high, they will feel uncomfortable (1). Moderate novelty will make them feel comfortable. In contrast, even if an experience is a novel experience, one gets used to it by experiencing it repeatedly. Therefore, if one is experiencing unpleasant novel events, one should get used to them earlier; if one is experiencing pleasant novel events, one should be as unaccustomed as possible. Understanding one’s response when repeatedly experiencing a novel event is important in maintaining long-term novelty. In the field of psychology, the attenuation of a response by repeatedly experiencing a stimulus is defined as habitation (2–6). Lécuyer (1989); this phenomenon suggested that the amplitude of a novelty reaction and habituation speed are linked to one’s attention and speed of information processing in the development stage (4). Croy et al. (2013) demonstrated that unpleasant stimuli initially caught more attention, and repeated exposure led to reduced emotional salience of unpleasant stimuli in the experiment using subjective evaluation and the event-related potential (ERP) (3). These studies indicate that habituation to novel stimuli is affected by attention.

Neuropharmacological studies using animals have investigated the neural mechanism underlying habituation to novelty (7–10). Habituation to novelty is explained by antagonistic modulation of the brain excitatory nervous system (acetylcholine, adrenaline, and glutamate) and inhibitory nervous system (gamma-aminobutyric acid). As Stein’s classic theory (6) states, a novel stimulus activates the excitatory mechanism, which in turn activates the inhibitory mechanism. The inhibitory mechanism becomes conditioned to the onset of the stimulus after repeated presentations; when a repeated presentation is predictable, conditioned activation of the inhibitory mechanism overrides the direct activation of the excitatory mechanism. This neuropharmacological theory shows that the predictability of novel stimuli plays an important role in the formation of habituation.

Our research group has developed a mathematical model of emotional dimension for novelty using information theory and a Bayesian approach (11, 12). The model formalizes arousal (primary emotional dimension) (13) as information gain (also called Kullback–Leibler divergence) obtained by experiencing events. Information gain is expressed as a function of predictability (i.e., uncertainty and prediction error). The functional model is supported experimentally with the gaze shift (14) and the event-related brain potential (ERP) (12) of human participants as indexes of arousal.

In this study, we assumed decrement of arousal evoked by the same event as habituation to novelty and aimed to elucidate how predictability affects the habituation to novelty. We investigated the effect of predictability on habituation to novelty by applying our mathematical model of arousal (12). We formulated habituation to novelty as a decrement in Kullback-Leibler divergence from Bayesian prior to posterior (information gain) representing arousal through Bayesian update. We then predicted the effects of the predictability (uncertainty and prediction error) on the time-course change of arousal as the primary factors constituting novelty by mathematical simulation of the model. We confirmed the biological validity of the model by experimentally demonstrating whether the model prediction and the activity in the human brain are consistent. Polich et al. investigated the brain activity related to habituation to stimuli using P300, which is one component of ERP, and reported the characteristics of attenuation of brain activity by repeated stimulus presentations (15–20). P300 (also known as P3) is a positive component that appears at a latency of approximately 250 to 600 milliseconds among the components of ERP (21). P300 for stimulus is known to predominantly appear from the frontal to the central and parietal brain parts and is considered to reflect attention to stimuli. In recent years, some studies have investigated habituation to stimuli using P300 (22–25), but the effects of predictability (uncertainties and prediction errors) on brain activity related to novelty habituation remain unclear. In this study, we demonstrated an experimental evidence of the model prediction using P300. We used the experimental task employed in our previous study (12) to examine the brain response to novelty.

## 2 Modeling habituation to novelty

We mathematically formulated habituation to repeated exposure of novel stimuli based on our previously proposed model of emotional dimensions associated with novelty (12, 26, 27).

A novel event provides new information. We used the amount of information acquired by an event as the extent of novelty. Considering a transition before and after experiencing an event, we assumed a Bayesian update of one’s belief from prior to posterior. We defined the amount of information gained from the event as Kullback–Leibler divergence from prior to posterior, which we termed *information gain*. Information gain is a decrease in self-information averaged over posterior. Accordingly, the information gain represents a decrease in uncertainty by experiencing an event. In addition, the information gain also represents surprise (14) and emotional arousal (12). When one repeatedly experiences the same event, uncertainty and surprise (i.e., information gain) to the event should decrease. We, therefore, assumed that the decrement in information gain represents habituation to a novel event.

### 2.1 Bayesian update model

Our Bayesian model assumed that one estimates a parameter *θ* using both one’s prior *p*(*θ*) and continuous data *x* ∈ *R* obtained by experiencing an event (12). The Bayes’ theorem updates the prior to the posterior *p*(*θ*│*x*) as the following equations:

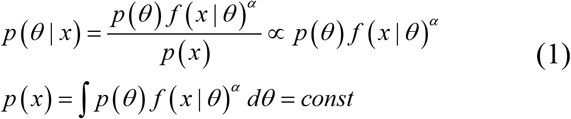

Posterior is proportional to a product of prior and a likelihood function *f*(*x*|*θ*) because the denominator *p*(*x*), or evidence, is constant. α is termed *learning rate* (28) that adjusts the amount of the prior update.

Assuming that one experiences the identical event and obtains the same data (*x*) *k* times, the *k*th posterior *p*_*k*_(*θ*│*x*) is proportional to a product of the initial prior and the likelihood functions when the likelihood functions are independent distributions:

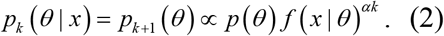

where the *k*th posterior is used as *k*+1th prior *p*_*k* + 1_(*θ*). Assuming that one’s brain encodes *n* samples of the identical data *x* as a Gaussian distribution *N*(*μ*,*σ*^2^) with a flat prior, using the distribution as likelihood function and the formula (2), a nonflat prior of *μ* following a Gaussian distribution *N*(*η*,*τ*^2^) is updated to the following Gaussian distributions:

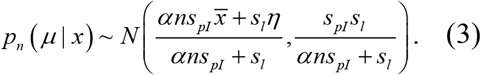

where 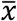 is the mean of the data, *s*_*pl*_ = *τ*^2^, and *s*_*l*_ = *σ*^2^.

### 2.2 Arousal update model (habituation)

Information gain (*G*_*n*_) in the *n*th repeated exposure of the identical continuous data or stimulus *x* is written as Kullback–Leibler divergence from posterior to prior:

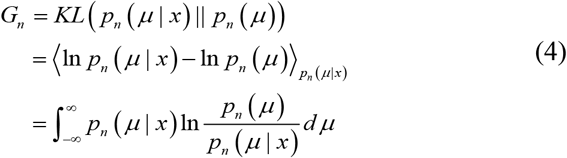

With the assumption of the Bayesian update in formula (2), we replace the *n*th prior by *n-1*th posterior.

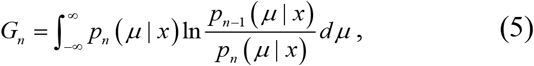

When the posterior follows the Gaussian posterior of formula (3), we derive the *n*th information gain as a function of initial parameters:

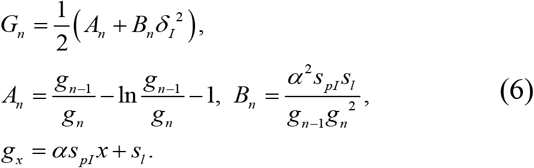

We term 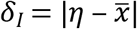 as the *initial prediction error* that represents the absolute difference between the prior mean and peak of the likelihood function. We term the variance of prior *s*_*pl*_ as *initial uncertainty*. The variance of the data *s*_*l*_ refers to *external noise* in the case of sensory data (i.e., stimuli). From formula (6), information gain is a function of the following three parameters: initial prediction error, initial uncertainty, and external noise.

### 2.3 Effects of initial prediction errors and initial uncertainties on the habituation to novelty

We analyzed how initial uncertainty and initial prediction error affect the decay of information gain or habituation. Figure 1 shows the decay of information gain as a function of the number of update by repeated exposures to the same data for varied initial prediction errors when the initial uncertainty is fixed. The information gain increases with the initial prediction error at any number of update *n* ∈ ℕ.

**Figure 1.**
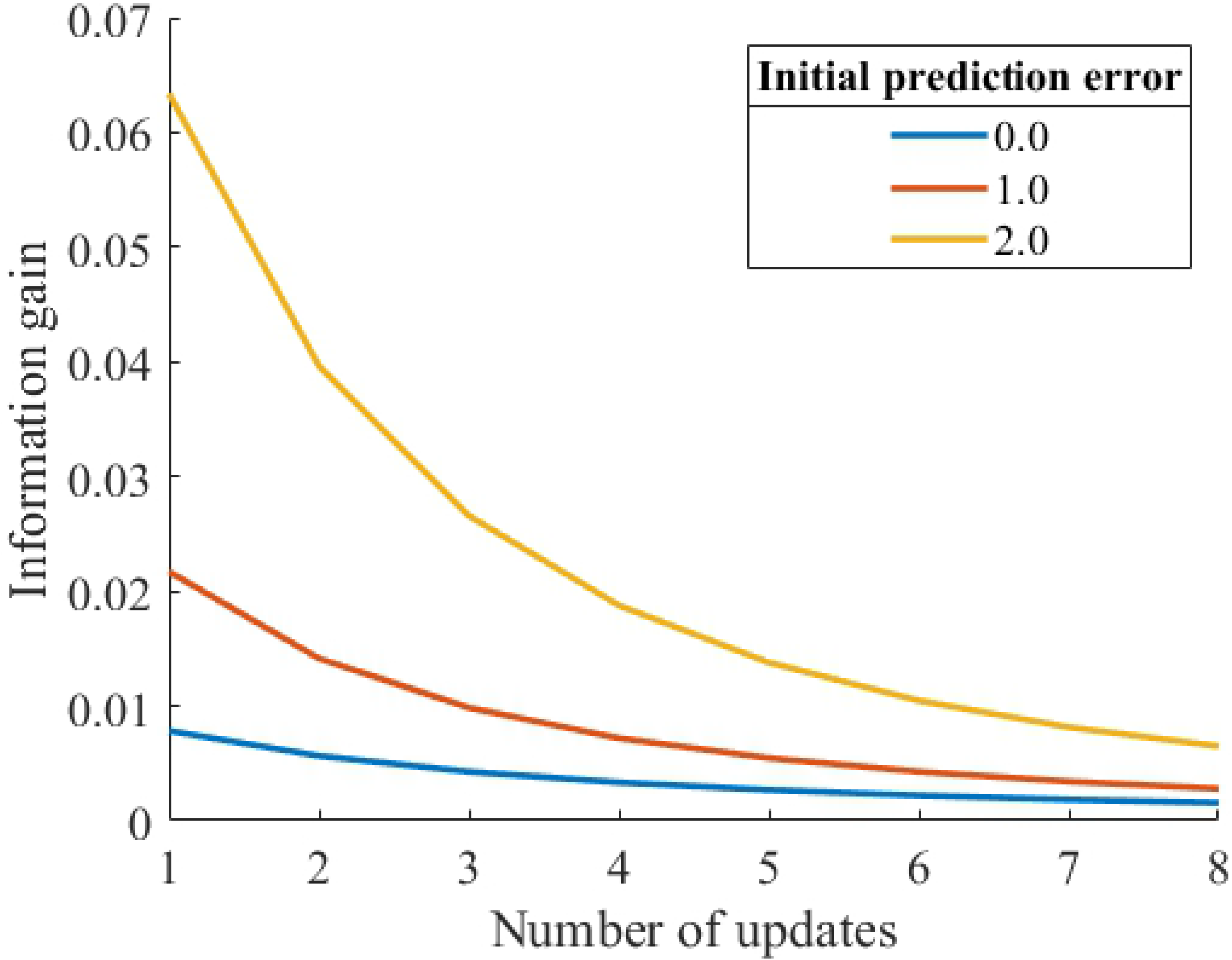
Updates of information gain for different initial prediction errors (initial uncertainty = 1.0, noise = 0.5, learning rate *α* = 0.1)

Figures 2 and 3 show the decay of information gain with the number of updates for different initial prediction errors (0.0 and 10.0). When the initial prediction error is 0.0, the larger initial uncertainties result in larger information gains. By contrast, when the initial prediction error is 10.0, larger initial uncertainties result in smaller information gain. That is, the effect of uncertainty on information gain is reversed for these two different prediction errors. In the case of n=1 update, this reversion occurs when the relationship between different initial uncertainties *s*_*p*1_ and *s*_*p*2_ is as follows (12):

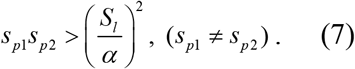

**Figure 2.**
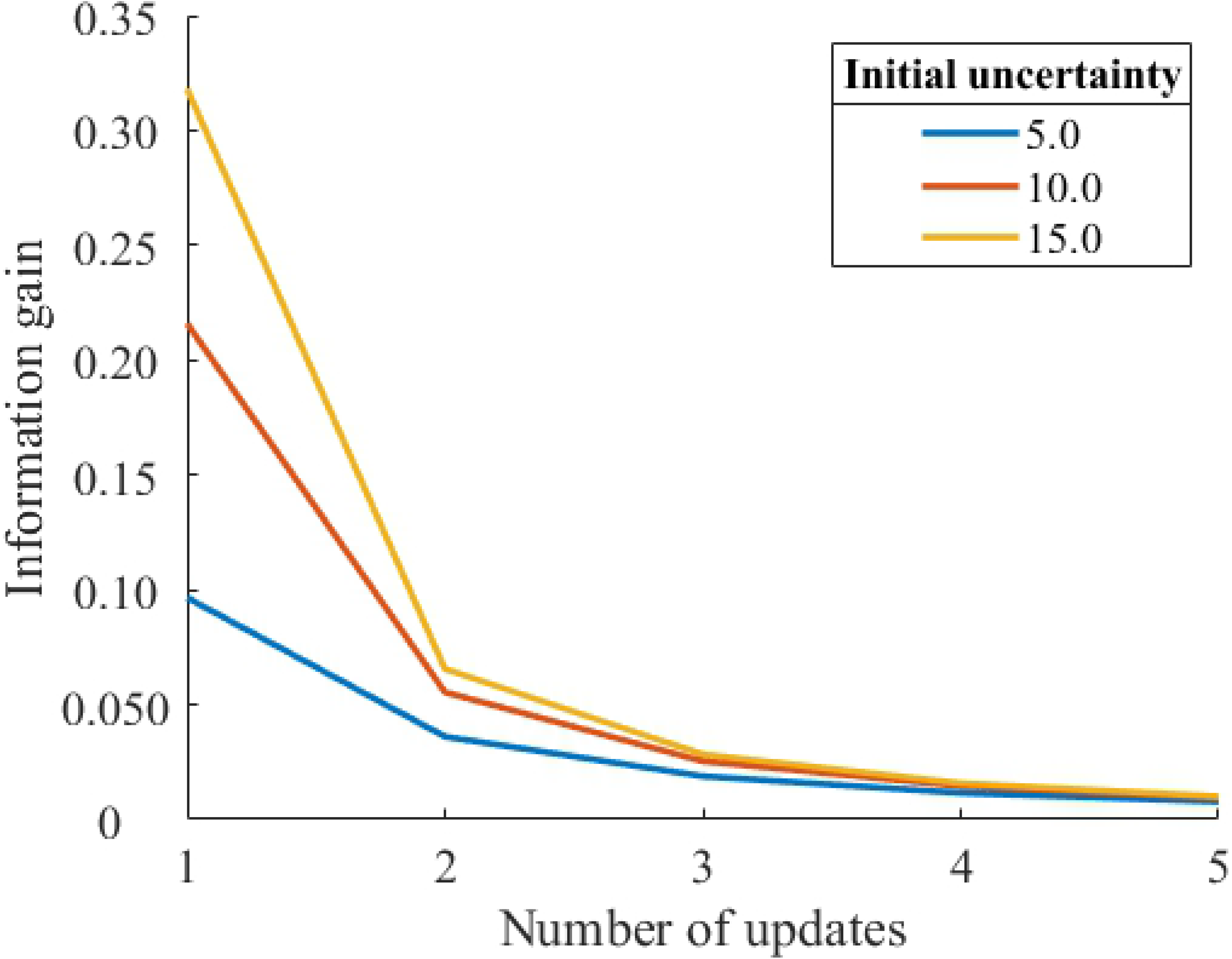
Updates of information gain for different initial uncertainties (initial prediction error = 0, noise = 0.5, learning rate *α* = 0.1)

**Figure 3.**
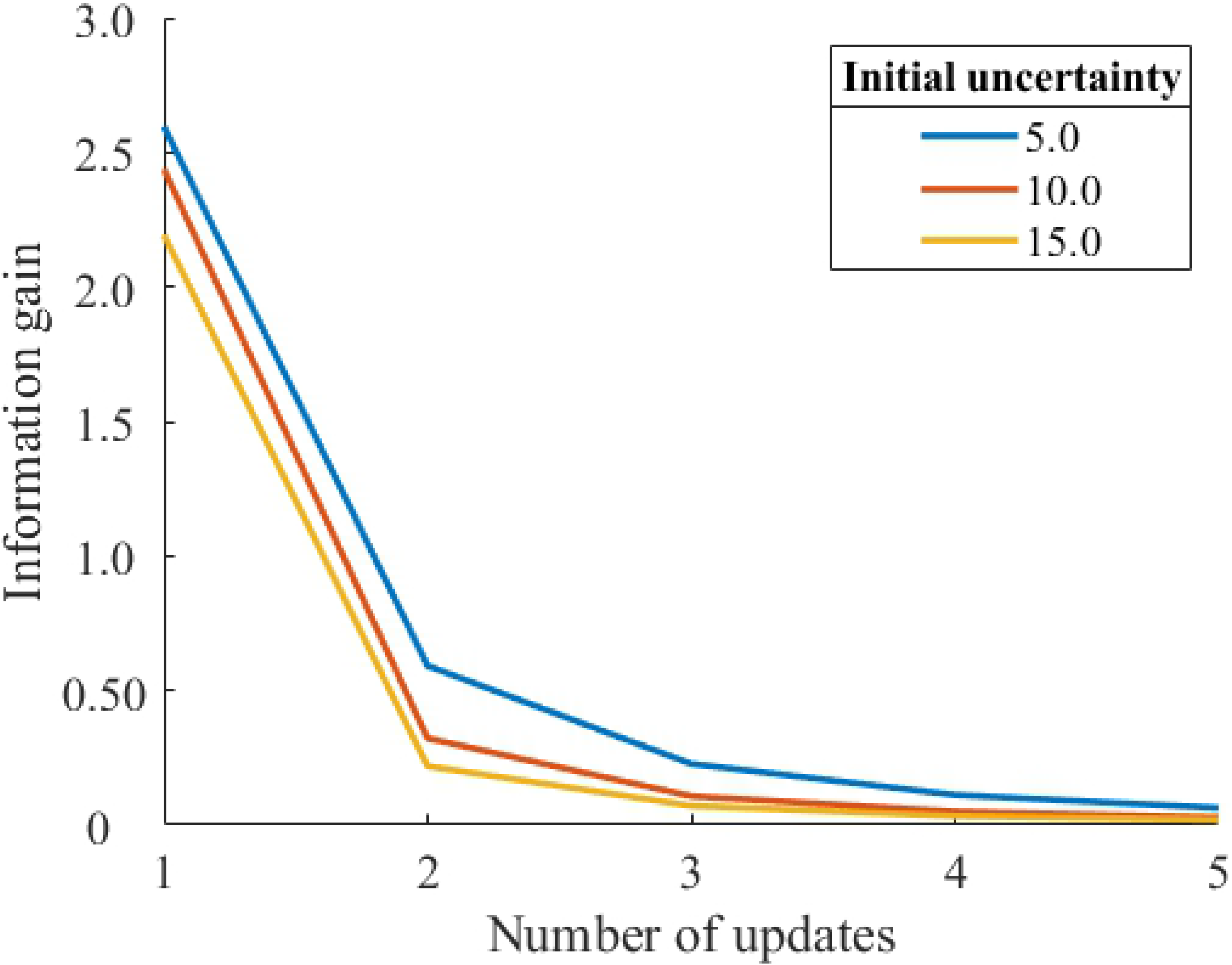
Updates of information gain for different initial uncertainties (initial prediction error = 10.0, noise = 0.5, learning rate *α* = 0.1)

As shown in both Figures 2 and 3, a larger initial uncertainty decreases the information gain more significantly from *n* = 1 to *n* = 2. As shown in Fig. 2, the larger information gain with a larger initial uncertainty quickly decreases to the same level of smaller initial uncertainty conditions. As shown in Fig. 3, the information gain with a lager initial uncertainty converges to zero faster. These simulation results imply that a greater initial uncertainty tends to result in a faster decay of information gain by updating, suggesting that lager initial uncertainty results in faster habituation.

Integrating information gain G with the number of updates n gives the following:

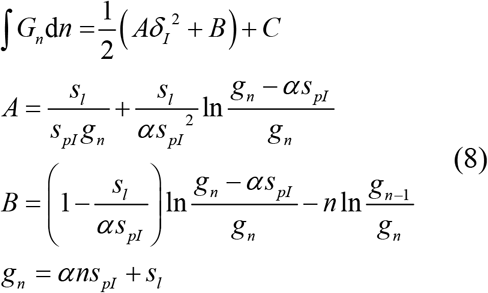

C is the integration constant. Substituting infinity for n of the above indefinite integral gives:

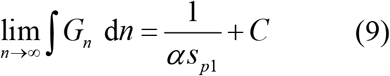

Equation (7) shows that the larger the value of the initial uncertainty, the smaller the sum of the information gain obtained when the stimulation is repeated indefinitely. As shown in Figure 3, the larger the value of the initial uncertainty, the smaller the initial value of the information gain. The analysis shows that this relationship remains even if n is increased to infinity. In other words, for any number of updates, the larger the value of the initial uncertainty, the smaller the information gain. Figure 4 shows that the value of the information gain when n is infinite becomes smaller as the value of the initial uncertainty is larger, regardless of the value of the initial prediction error.

**Figure 4.**
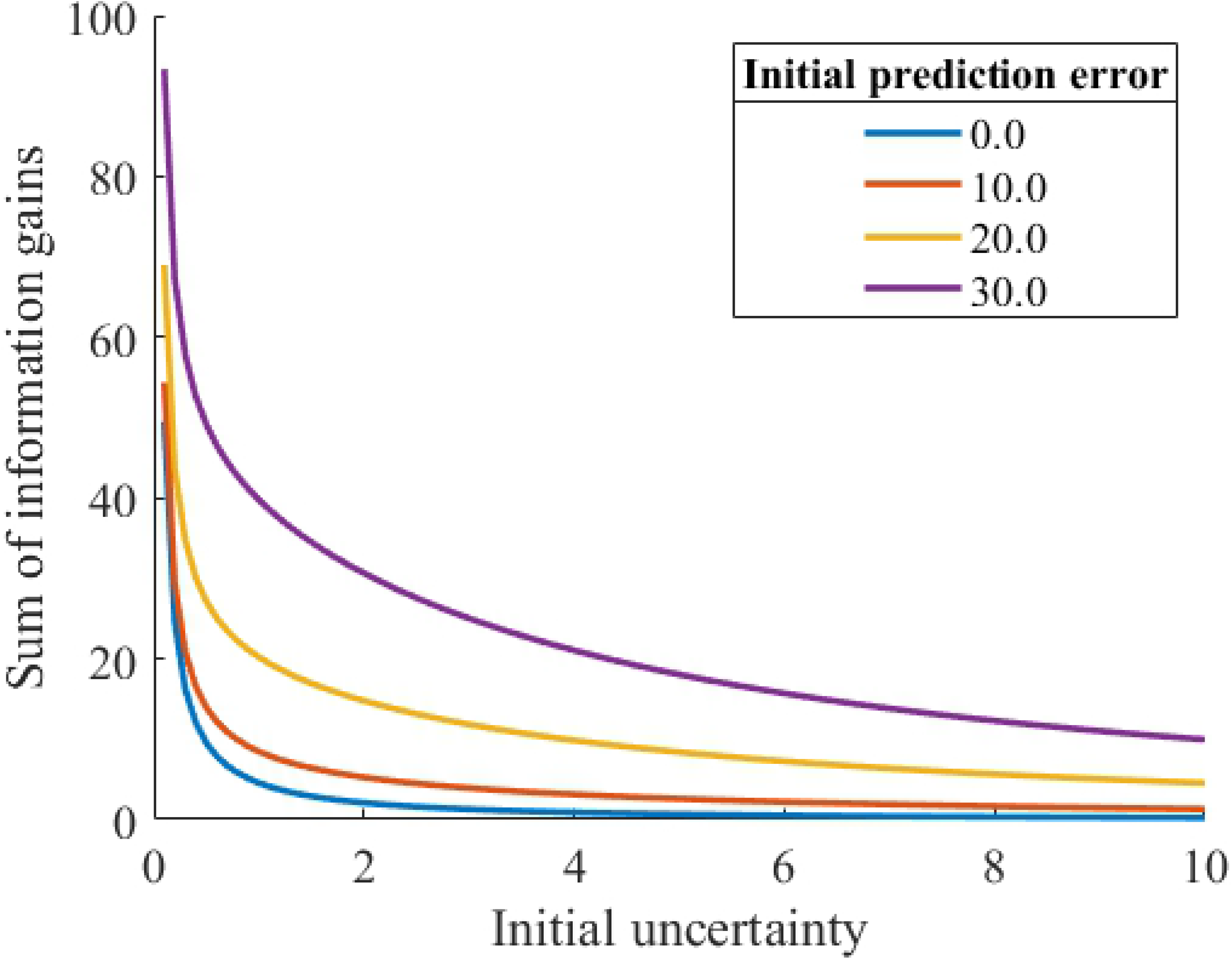
Relationship between the initial uncertainty and the integrated value of the information gain

## 3 Experiment

We conducted an experiment using electroencephalogram (EEG) recordings and questionnaires to test our hypotheses derived from the mathematical model predictions. We tested the hypothesis (H1) that the larger the initial uncertainties, the faster the surprise decays, regardless of the initial prediction error. In addition, we tested the hypothesis (H2) that larger initial uncertainties result in larger surprises when the initial prediction errors are small and smaller surprises when the initial prediction errors are large. We quantified habituation of surprise intensity using a four-level Likert scale and P300 amplitudes (12, 29). This experiment was based on the methodology in our previous study (12).

### 3.1 Materials and Methods

#### 3.1.1 Participants

Participants were eight right-handed adult males (age range: 21–27 years) who had no brain-related disorders, abnormalities associated with their eyesight, or other diseases. Handedness was assessed by the FLANDERS handedness questionnaire (30). This study was approved by the Research Ethics Committee of the University of Tokyo, Graduate School of Engineering. All participants gave their consent to participate in this study.

#### 3.1.2 Stimuli

We used four types of short video stimuli (duration: 2,500 ms) in which a percussion instrument was struck once and a synthesized percussive sound followed (see our previous study for details) (12). We performed an experimental manipulation of initial uncertainties due to the familiarity of percussion instruments. The clave and hand drum were used as familiar percussion instruments (i.e., low initial uncertainty, A; Table 1), and the jawbone and slit drum were used as unfamiliar percussion instruments (i.e., high initial uncertainty, B; Table 1). We manipulated the initial prediction errors by the degree of congruency between the percussion instrument and the synthesized percussive sound. We used synthesized percussive sounds that were consistent with the instruments shown in congruent conditions (i.e., low initial prediction error, X; Table 1). In contrast, we used sounds that were inconsistent with the instruments in incongruent conditions (i.e., high initial prediction error, Y; Table 1). For each video stimulus, a percussion instrument was first shown in the center of the screen; a percussion instrument was then struck once 500 ms after the onset of the video stimulus, and a percussive sound was presented simultaneously.

**Table 1.** Combination of percussion instruments and percussive sounds

|  | Instruments | Low initial prediction error | High initial prediction error |
| --- | --- | --- | --- |
|  |  | (Congruent sound, X) | (Incongruent sound, Y) |
| Low initial<br>uncertainty<br>(Familiar, A) | Clave | Clave (AX) | Bell (AY) |
|  | Hand drum | Hand drum (AX) | Guiro (AY) |
| High initial<br>uncertainty<br>(Unfamiliar, B) | Jawbone | Jawbone (BX) | Vibraphone (BY) |
|  | Slit drum | Slit drum (BX) | Snare (BY) |

#### 3.1.3 Procedure

In the experiment, participants watched video stimuli while EEG recordings were taken and answered subjective feelings of surprise in an electromagnetically shielded room. The experiment consisted of 480 trials (eight videos [Table 1] × 60 presentation sets). The inter-trial interval was 1,000–2,000 ms. The eight videos were presented in a random order in each set. Participants reported the intensities of their surprise using a four-level Likert scale upon listening to the percussive sounds during the first, 20th, 40th, and final presentation sets.

#### 3.1.4 EEG Measurement

We recorded EEGs during experimental tasks using an EEG amplifier system (eego sports, ANT Neuro) with active electrodes (sampling rate: 1000 Hz, time constant: 3 s). EEGs were recorded from electrodes positioned at the Fz, Cz, and Pz points according to the international 10–20 system (31) with reference to the nose. To monitor the blinking of the eye, Fp1 point was recorded. All electrode impedances were below 60 kΩ.

#### 3.1.5 Analysis

Averaged ERP waveforms were computed from 200 ms before the video stimulus onset (i.e., the start of the video) to 1,500 ms after the video stimulus onset following the application of a digital band-pass filter of 0.1–20 Hz. Waveforms were aligned to the 200 ms pre-stimulus baseline period. The averaging was performed for each participant, stimulus type (i.e., AX, AY, BX, and BY), and the number of exposure (i.e., 1-40, 41-80, and 81-120) for video stimuli. To calculate the average waveforms of P300 for each participant, each number of exposure consisted of 40 trials. Data of one participant were excluded from the ERP analysis due to excessive eye-blink artifacts. Ocular artifacts (eye movements and blinks) and muscle artifacts were removed by the Automatic Subspace Reconstruction method (32). Any epochs containing EEG signals exceeding ± 80 μV were regarded as artifacts and were removed. P300 was defined as the largest positive peak occurring 250–600 ms after the onset of the percussive sound. The baseline-to-peak amplitude of P300 was measured at the Pz point to examine the parietal P300, which represents surprise (12, 29). Repeated-measures analysis of variance (ANOVA) was used to analyze the Likert scale and P300 amplitudes data. Statistical significance was defined as p < 0.05.

## 4 Results

Figures 5 and 6 show the average subjective scores of surprise for the number of exposure in congruent and incongruent conditions (i.e., low and high initial prediction errors, respectively), respectively. In both lower and higher initial uncertainties, the first exposure had the largest score, and subsequent exposures decreased the score. A three-factor repeated-measures ANOVA was performed with the initial uncertainty, the initial prediction error, and the number of exposure as independent variables and the score of surprise as a dependent variable. The main effects were significant for the initial prediction error (*F* = 8.948, *p* = .020) and the number of exposure (*F* = 15.598, *p* = .000). The score of surprise for the high initial prediction error was large than that for the low initial prediction error. In addition, the higher the number of exposure, the smaller the score of surprise. The interaction effect of the initial uncertainty and the initial prediction error was significant (*F* = 11.324, *p* = .012). The simple main effect of the initial uncertainty was significant for both the low initial prediction error (*F* = 9.289, *p* = .019) and the high initial prediction error (*F* = 9.215, *p* = .019). The score of surprise under the high initial uncertainty was greater than that under the low initial uncertainty when the initial prediction error is low. In contrast, the score of surprise under the low initial uncertainty was greater than that under the high initial uncertainty when the initial prediction error is high. The reversal of subjective surprise for the initial uncertainty due to the initial prediction errors might reflect the simulation results, as shown in Figures 2 and 3. These results supported the hypothesis H2.

**Figure 5.**
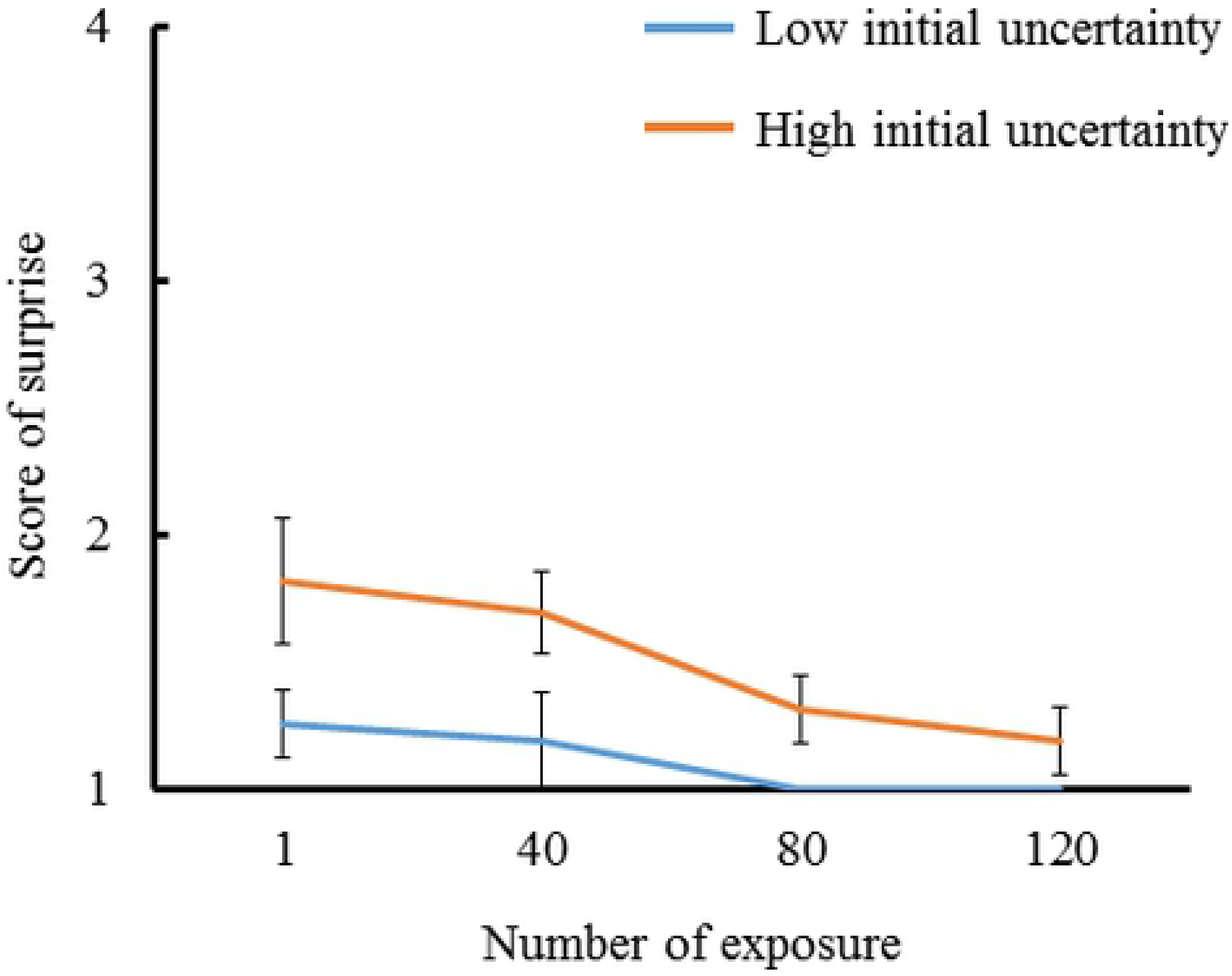
Subjectively reported scores for surprise intensities in response to percussive sounds congruent with the instrument shown (i.e., low initial prediction error). The results for familiar and instruments were compared at every 40 exposures (N = 8).

**Figure 6.**
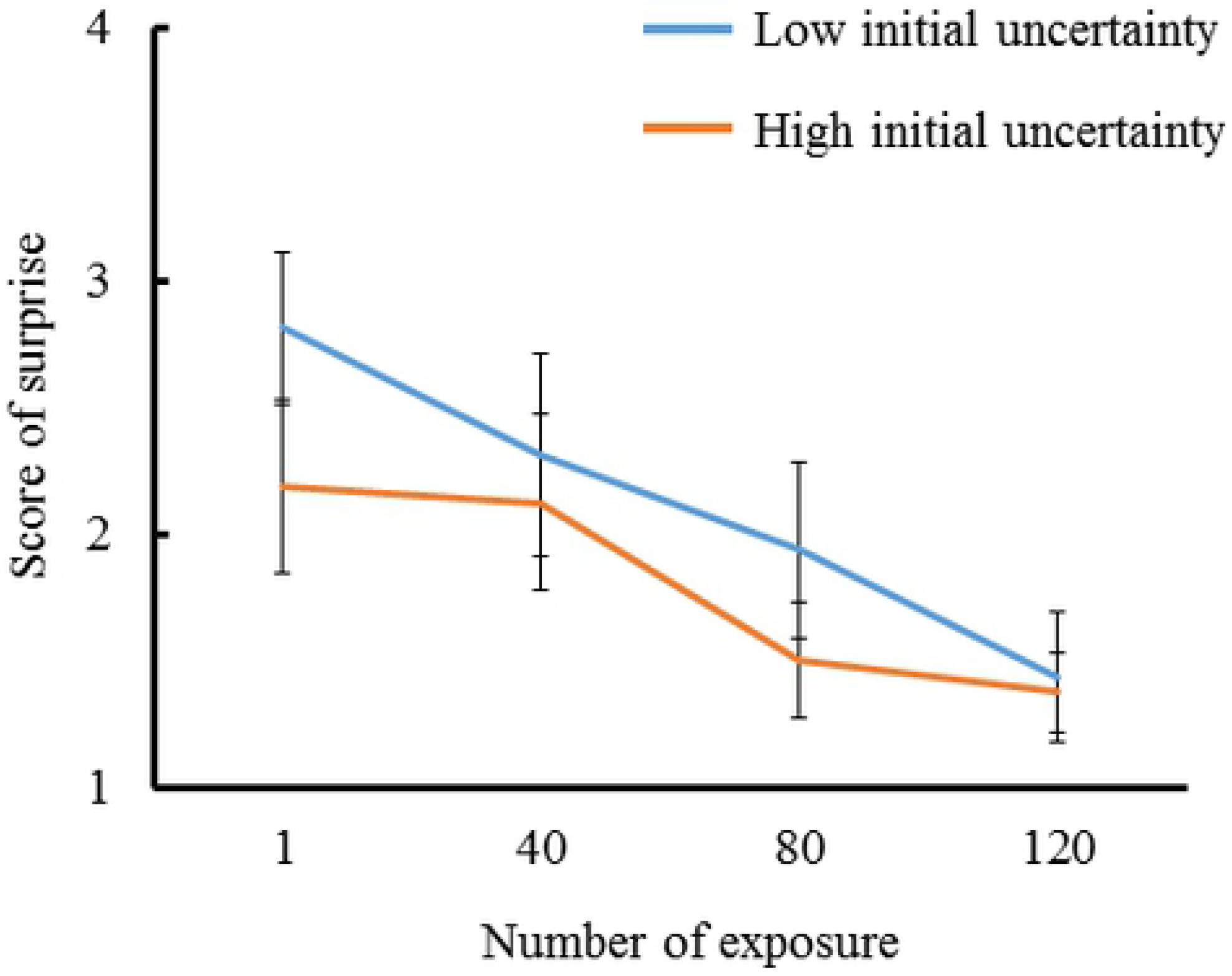
Subjectively reported scores for surprise intensities in response to percussive sounds incongruent with the instrument shown (i.e., high initial prediction error). The results for familiar and unfamiliar instruments were compared at every 40 exposures (N = 8).

Figure 7 shows the grand mean ERP waveforms for the number of exposure in the four congruent and incongruent conditions (i.e., low and high initial prediction errors, respectively), respectively. In both low and high initial prediction errors, the first 40 exposures had the largest P300 amplitude, and subsequent exposures attenuated the amplitude in the unfamiliar condition (i.e., high initial uncertainty). On the other hand, in the familiar condition (i.e., low initial uncertainty), the P300 amplitude gradually decreased with the increasing number of exposure in the low initial prediction error, and the P300 amplitude was maintained at the same level as that in 40 and 80 exposures and then decreased in the high initial prediction error.

**Figure 7.**
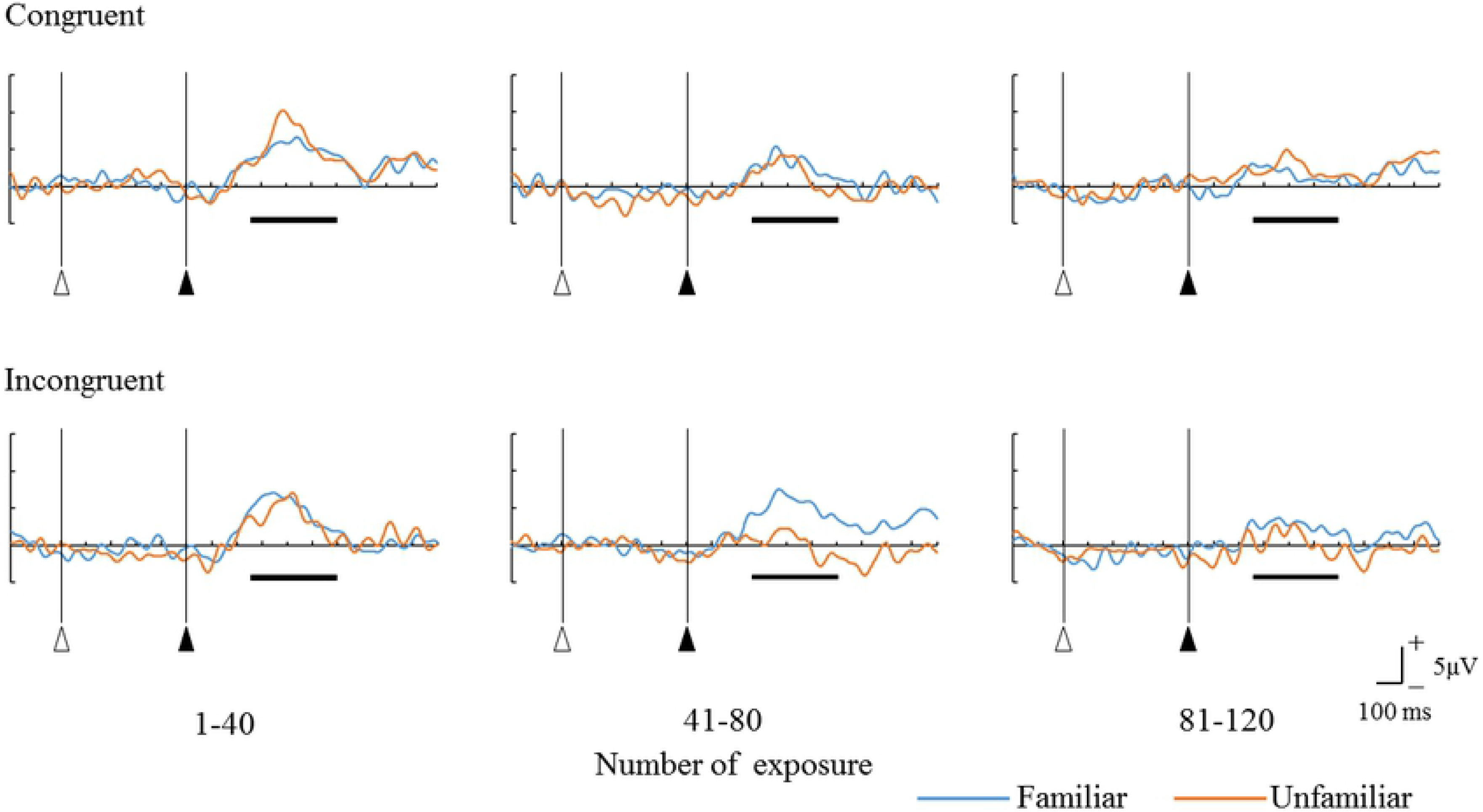
Grand mean ERP waveforms for the four combinations of percussion instruments and percussive sounds at the parietal midline region (Pz). Open triangles: the onset of film presentation. Solid triangle: the onset of beating sound presentation. The horizontal bars show the time range of 250 – 600 ms for the P300 latency.

Figures 8 and 9 show the average P300 amplitude for the number of exposures in congruent and incongruent conditions (i.e., low and high initial prediction errors, respectively). The P300 amplitude for the high initial uncertainty was larger than that for the low initial uncertainty in the low initial prediction error, and the P300 amplitude for the low initial uncertainty was larger than that for the high initial uncertainty in the high initial prediction error. A three-factor repeated-measures ANOVA was performed with the initial uncertainty, the initial prediction error, and the number of exposures as independent variables and the P300 amplitude as a dependent variable. The main effect of the number of exposure was significant (*F* = 8.447, *p* = .005). The P300 amplitude for 81–120 exposures was larger than that for 1–40 and 41–80 exposures. The interaction effect of the initial uncertainty and the number of exposures was significant (*F* = 4.114, *p* = .044). The simple main effect of the number of exposures was significant for both the low initial uncertainty (*F* = 6.046, *p* = .015) and the high initial uncertainty (*F* = 8.697, *p* = .005). The P300 amplitude for 81–120 exposures was smaller than that for 41–80 exposure when the initial uncertainty is low. In contrast, the P300 amplitudes for the 41–80 and 81– 120 exposures were smaller than those for 1–41 exposures when the initial uncertainty is high. Therefore, the larger the initial uncertainty, the sooner the P300 decay (that is, the faster the reduction of surprise). This tendency of the 300 amplitude decay is consistent with the results of our model predictions shown in Figures 2 and 3. These results supported the hypothesis H1.

**Figure 8.**
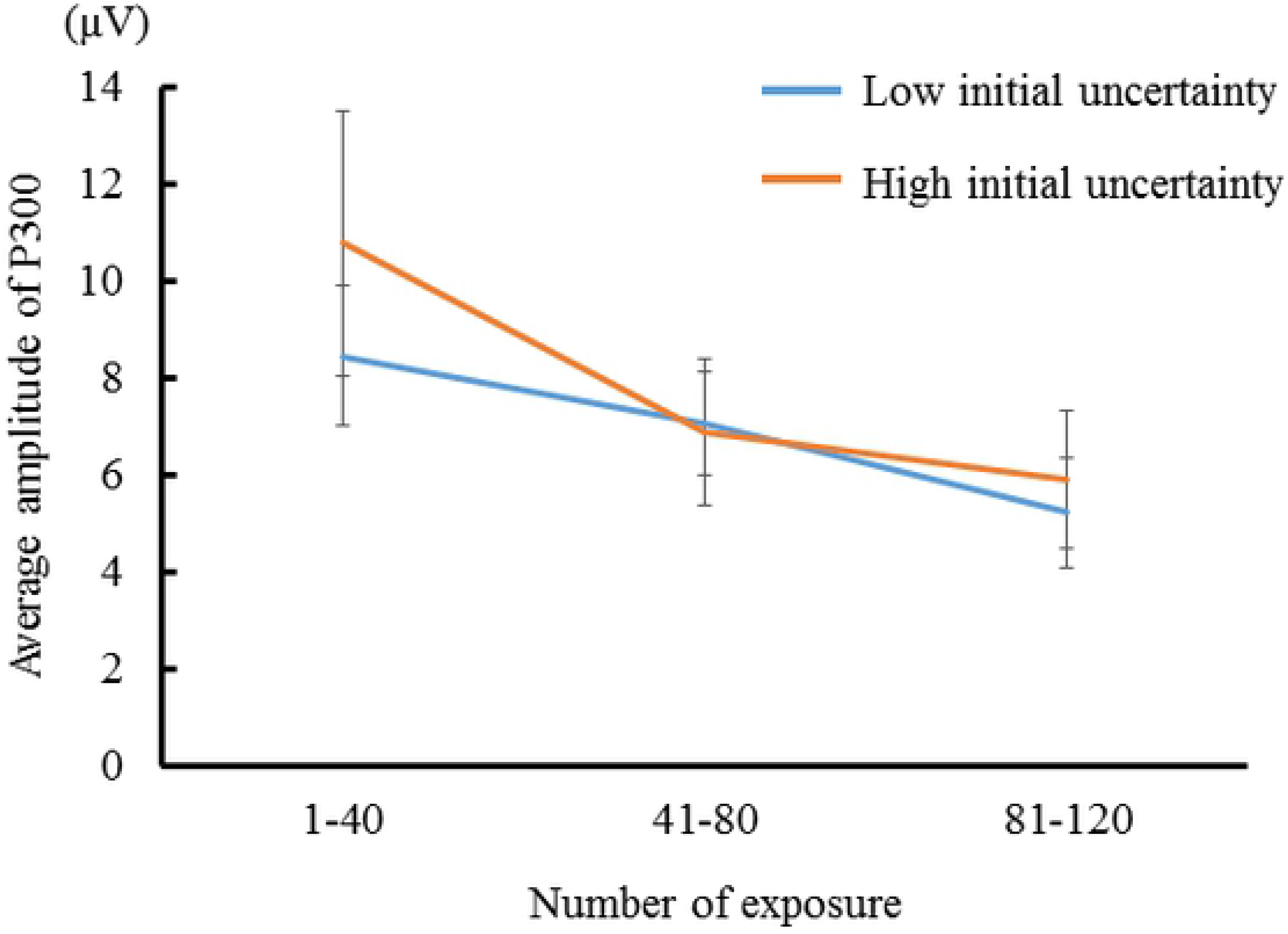
P300 amplitudes evoked by percussive sounds congruent with the instrument shown (i.e., low initial prediction error). The results for familiar and unfamiliar instruments were compared at every 40 exposures (N = 7).

**Figure 9.**
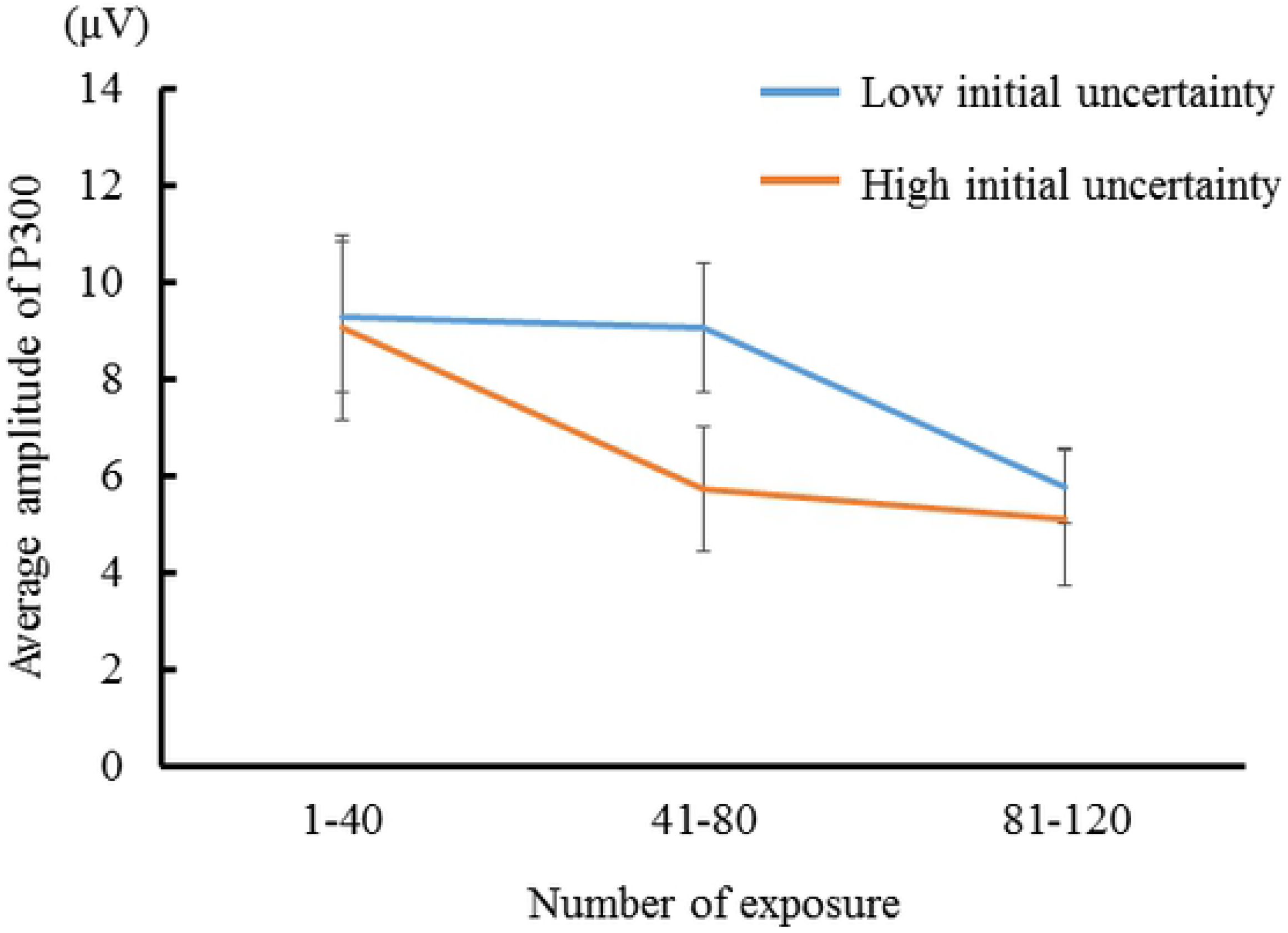
P300 amplitudes evoked by percussive sounds incongruent with the instrument shown (i.e., high initial prediction error). The results for familiar and unfamiliar instruments were compared at every 40 exposures (N = 7).

## 5 Discussion

In this study, we predicted the effects of predictability on habituation to novelty by applying the Bayesian update model. In our model, we assumed a decrement in information gain (i.e., arousal) as habituation to novelty. We then conducted an event-related potential experiment to demonstrate an experimental evidence of the model prediction. The model formalized habituation as a decrement in information gain (i.e., decay of surprise). We formalized the information gain as Kullback–Leibler divergence from prior to posterior based on Bayesian update. Based on this model, posterior is proportional to a product of prior and likelihood function. With the Gaussian prior and likelihood function, we derived the information gain as a function of three parameters: initial uncertainty, initial prediction error, and noise of sensory stimulus.

We found an interaction effect between initial uncertainty and initial prediction error on habituation expressed as decrement in information gain based on mathematical simulation in the experiment using P300 and questionnaires, and the findings indicate that the greater the initial uncertainty, the faster the information gain decreases and converges to zero. As previous studies (12, 33) demonstrated, the initial uncertainty depends on one’s prior knowledge and experience. More prior knowledge and experience result in less uncertainty. We assumed that the affinity that comes from familiarity of the object decreases the uncertainty. We conducted a P300 experiment using a set of videos of percussion instruments accompanied by synthesized percussive sounds. We manipulated the initial uncertainty with the familiarity of instruments shown and the initial prediction error with the congruency of percussive sounds. We used P300 amplitudes as an index of how the participants are surprised by the percussive sounds (12). The experimental results of P300 amplitudes support the hypothesis: the less familiar the object, the faster one becomes accustomed to novel stimuli. Consistent with the simulation results, when the uncertainty was high, the degree of information gain was changed greatly in the time transition from the initial exposure.

Brain activity related to habituation to stimuli has been investigated in many studies, including a series of studies by Polich et al. (15–20, 22–25). This present study clarified for the first time the influence of uncertainty and prediction error on brain activity related to habituation to novel stimuli based on mathematical models. The results of this study may indicate that when attention is paid more strongly to novel stimuli (e.g., high uncertainty situation), the initial information gain increases, and accordingly, information processing is promoted, resulting in rapid habituation. In addition, Lécuyer (1989) stated that the amplitude of novelty reaction and habituation speed are linked to one’s attention and speed of information processing (4), and a neuropharmacological study pointed out the relationship between predictability and habituation of novel stimuli (6). Our results suggest that in highly uncertain situations, repeated exposure to stimuli may increase predictability and enhance habituation to novel stimuli.

This study investigated the effects of initial uncertainty and prediction error on the habituation (decrease of arousal level) for novelty based on our mathematical models and psychophysiological experiments. We introduced the concept of Bayesian update and formulated the mechanism by defining the habituation with novelty as a decrease in information gain. The results of the simulations and experiments in this study suggest the effect of initial uncertainty on the degree of surprising attenuation for novelty and the interaction due to prediction errors. Uncertainties in this model include parameters that correspond to individual knowledge, experience, frequency of contact with events, familiarity, and typicality. Uncertainty is a factor that can explain variations in the habituation to novelty due to individual and subjective attributes. Because this study was conducted using a within-participant factorial design, the impact of individual differences on the results between conditions was small. However, we believe it is necessary to increase the number of participants in future experiments. Although we experimentally examined the effect of the familiarity of events on the habituation of novelty, we also need to experimentally test other parameters of uncertainty in the future work.

## Acknowledgements

We thank Prof. Tamotsu Murakami and the members of the Design Engineering Laboratory at the University of Tokyo for supporting this project.

